# An in-vivo validation of ESI methods with focal sources

**DOI:** 10.1101/2021.09.10.459782

**Authors:** Annalisa Pascarella, Ezequiel Mikulan, Federica Sciacchitano, Simone Sarasso, Annalisa Rubino, Ivana Sartori, Francesco Cardinale, Flavia Zauli, Pietro Avanzini, Lino Nobili, Andrea Pigorini, Alberto Sorrentino

**Affiliations:** Department of Mathematics, Università degli Studi di Genova, Genoa, Italy; CNR - SPIN, Genoa, Italy; CNR - IAC, Rome, Italy; Department of Biomedical and Clinical Sciences “L. Sacco”, Università degli Studi di Milano, Milan, Italy; Department of Neurosciences, Center for Epilepsy Surgery “C. Munari”, Hospital Niguarda, Milan, Italy; CNR - Istituto di Neuroscienze, Parma, Italy; Child Neuropsychiatry Unit, IRCCS “G. Gaslini” Institute, Genoa, Italy; DINOGMI, Università degli Studi di Genova, Genoa, Italy

**Keywords:** ESI, EEG, inverse methods

## Abstract

Electrical source imaging (ESI) aims at reconstructing the electrical brain activity from measurements of the electric field on the scalp. Even though the localization of single focal sources should be relatively straightforward, different methods provide diverse solutions due to the different underlying assumptions. Furthermore, their input parameter(s) further affects the solution provided by each method, making localization even more challenging. In addition, validations and comparisons are typically performed either on synthetic data or through post-operative outcomes, in both cases with considerable limitations.

We use an in-vivo high-density EEG dataset recorded during intracranial single pulse electrical stimulation, in which the true sources are substantially dipolar and their locations are known. We compare ten different ESI methods under multiple choices of input parameters, to assess the accuracy of the best reconstruction, as well as the impact of the parameters on the localization performance.

Best reconstructions often fall within 1 cm from the true source, with more accurate methods outperforming less accurate ones by 1 cm, on average. Expectedly, dipolar methods tend to outperform distributed methods. Sensitivity to input parameters varies widely between methods. Depth weighting played no role for three out of six methods implementing it. In terms of regularization parameters, for several distributed methods SNR=1 unexpectedly turned out to be the best choice among the tested ones.

Our data show similar levels of accuracy of ESI techniques when applied to “conventional” (32 channels) and dense (64, 128, 256 channels) EEG recordings.

Overall findings reinforce the importance that ESI may have in the clinical context, especially when applied to identify the surgical target in potential candidates for epilepsy surgery.

## 1. Introduction

Electrical source imaging (ESI) is a procedure that allows reconstructing the neural activity inside the brain from recordings of the electric potential, usually obtained at the scalp. ESI is a key element in multiple frameworks related to the analysis of EEG data, including identification of brain regions involved in specific tasks [1,2] and estimation of connectivity in task-related or spontaneous activity [3]. Moreover, recent evidence [4,5,6,7] suggests that it could be considered a valuable tool in the context of pre-surgical evaluation of epileptic patients; in this case accuracy and reliability become of paramount importance.

Despite its usefulness, the use of ESI in clinical practice is still rather complex, due to the presence of numerous subjective choices involved: first, the choice of the ESI method among the many available options; second, the choice of the input value(s) for the parameter(s) the method depends on, a choice often overlooked but of great importance.

From a mathematical perspective, it is well known that reconstruction of neural currents from EEG data is an ill-posed problem with no unique solution; uniqueness can be restored by inserting *a priori* information through a so-called *regularization* procedure. However, there is no general agreement about the quality and quantity of a priori information to be inserted, and hence multiple ESI methods bloom [8, 9]. In addition, regularization methods typically require knowledge of the SNR of the input data in the form of a *regularization parameter*, and some of them try to avoid the typical bias towards superficial sources by inserting an additional *depth weighting parameter*.

In practice, some of these choices may turn out to be useful as they help to overcome notorious limitations of currently available ESI methods, by allowing the user to exploit their prior knowledge; however, different choices eventually lead to different solutions, and it is often not obvious what the optimal choices are. On the other hand, there is increasing consensus that focal sources, such as those mostly involved in epilepsy, should be relatively easy to localize. To quote a recent book [10], “MEG data are usually not ambiguous; it is mostly obvious where the active areas are located”; and a similar concept is expressed for EEG in [9]. For this type of source, it seems appropriate to ask whether some methods are better than others, and what parameters should be used to obtain reliable and accurate localization.

As an additional difficulty, validation and comparison of ESI methods is itself a tricky task, as the true sources of experimental recordings are never known exactly. There are two typical workarounds to this problem. One is to renounce the complexity of experimental data and use synthetic data to assess the reconstruction error of ESI [11,12,13,14,15,16]; this approach can result in reasonable comparisons between different methods but can hardly be used to give realistic estimates of localization accuracy in experimental scenarios, as the data generation process, particularly the forward model, is necessarily simplified. The second possibility, used in a growing number of studies, is to evaluate the accuracy of localization through post-surgical outcomes in epileptic patients [16,17,18,19,20]; this approach overcomes the limitations of synthetic data but has its own drawbacks, including the fact that resolution is limited to the size of the resected area. In this study we overcome the limitations of both approaches by exploiting a recently published EEG dataset of scalp recordings for which the ground truth is known. The dataset has been obtained at Niguarda Hospital in Milan, Italy: it contains EEG recordings with 256 channels, collected during the presurgical evaluation of patients affected by drug resistant focal epilepsy. Specifically, High-Density Scalp EEG data were collected during Single Pulse Electrical Stimulation (SPES), that is a clinical procedure increasingly employed for brain mapping and for the identification of abnormal cortical excitability in patients with epilepsy [21, 22, 23, 24]. During SPES, a brief current pulse is injected between two adjacent leads, producing an electrical current whose location can be accurately determined. Since this electrical current is strong enough to produce a visible signal on scalp HD-EEG, the procedure generates experimental data of scalp potentials originating from precisely known locations inside the brain. The resulting dataset is characterized by a very high signal-to-noise ratio and is ideally suited to evaluate in vivo the performance of ESI [25] in the case of focal activity.

We compare ten different ESI methods, thus possibly providing the most extensive comparison thus far: we test dipole fitting, wMNE [26], LORETA [27], sLORETA [28], eLORETA [29], dSPM [30], RAP- MUSIC [31], Gamma Map [32], MxNE [33] and SESAME [34].

For each method under consideration, we test several values for each input parameter, so as to verify (i) what is the optimal reconstruction attainable by an expert user who is capable of setting the parameter values correctly and (ii) to what extent the method is tolerant with respect to mis-specifications of these values. We remark that, in general, it is not straightforward what values to use in practice and in many recent studies this information is not present. In view of recent efforts to set up best practice guidelines of describing EEG/MEG studies [35], reporting the values of these input parameters is of fundamental importance to ensure reproducibility and replicability of the study. At the same time, a certain degree of tolerance with respect to the input parameter is a desirable property of ESI methods, as it removes part of the subjectivity in the analysis.

In summary, based on a unique dataset in which the sources of the EEG activity within the brain are known - i.e. a ground truth for the inverse solution methods is available - we compared the performances of the most commonly used ESI methods and, for each method we optimized the input parameters. The final aim is to provide a validation and a comparison of ESI methods based on a ground truth

## 2. Methods

### 2.1 Description of the data

The dataset used in this study is publicly available and has been described in [25]; in the following subsections we briefly summarize the main relevant features.

#### 2.1.1. Electrical Stimulation

Subjects had implanted intracranial shafts for pre-surgical evaluation of epilepsy; electrode positions were therefore established based on clinical needs, and ranged for each subject from superficial to deep locations: the distance from the closest sensor ranged from 28 to 64 mm. Electrical currents were delivered through platinum-iridium semi flexible multi-contact intracerebral electrodes (diameter: 0.8 mm; contact length: 2 mm, inter-contact distance: 1.5 mm; Dixi Medical, Besancon, France). Currents lasted 0.5 ms, were repeated either every 2 s (for 1 mA and 5 mA) or every 1 s (otherwise), and had intensities ranging between 0.1 mA and 5 mA. The number of recorded trials was either 40 (for 1 mA and 5 mA) or 60 (otherwise).

Electrode positions were measured by co-registering the post-implant CT (O-arm 1000 system, Medtronic) to the pre-implant MRI by means of the FLIRT software [36]. The location of every single lead was assessed using Freesurfer [37], 3D Slicer [38] and SEEG assistant [39]. When the EEG digitization MRI was different from the pre-implant MRI, transformation of the SEEG space to the EEG space was performed using an affine transformation between MRIs calculated with the ANTs software [40]. Normalized contacts’ coordinates were estimated through a non-linear registration between the subject’s skull-stripped MRI and the skull-stripped MNI152 template [41], using ANTs’ SyN algorithm. The accuracy of the normalization procedure was verified by visual inspection.

#### 2.1.2. High Density EEG recordings

256 EEG channels (Geodesic Sensor Net; HydroCel CleanLeads) were recorded with an EGI NA-400 amplifier (Electrical Geodesics, Inc; Oregon, USA) at a sampling frequency of 8,000 Hz, using a custom-made acquisition software, based on EGI’s AmpServerPro SDK and written in C++ and Matlab. No software filters were used during acquisition. The location of EEG electrodes and anatomical fiducials were digitized with a SofTaxicOptic system (EMS s.r.l., Bologna, Italy), coregistered with a pre-implant MRI (Achieva 1.5 T, Philips Healthcare).

#### 2.1.3. Generation of evoked responses

Raw data were high-pass filtered at 0.1 Hz (FIR filter; zero phase; Hamming window; automatic selection of length and bandwidth); for two subjects, data were also notch filtered at 50, 100, 150 and 200 Hz due to the presence of line noise. After rejection of bad channels through visual inspection, epochs were generated from −300 ms to 50 ms with respect to the electrical stimulation.

Evoked potentials were generated by averaging across all epochs produced by stimulation of the same contact pair. This produced very clear dipolar patterns produced by a single source, with a high Signal-to-Noise Ratio (SNR). In Figure 1 we report an example of a butterfly plot in the time window [−0.5; 1]ms.

**Figure 1:**
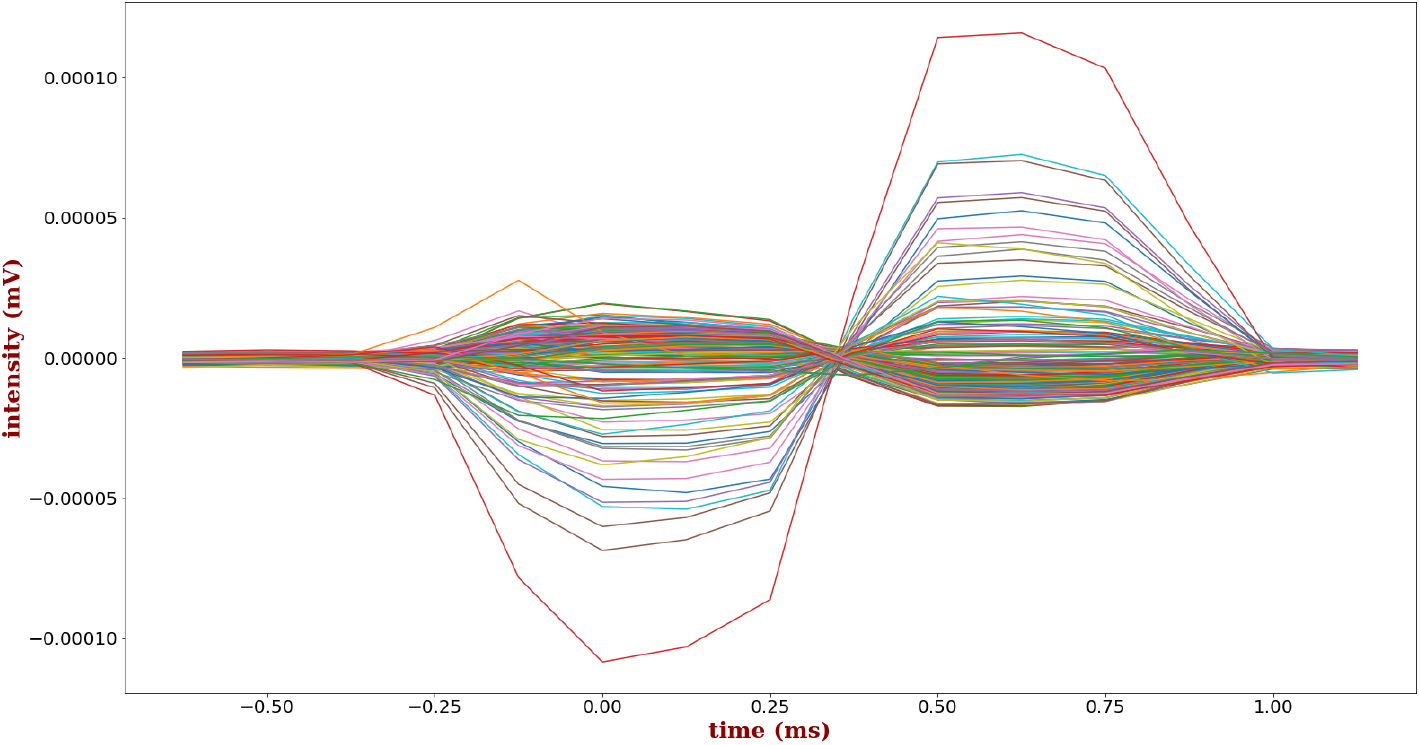
An example of evoked potential

Overall, the dataset analyzed in this study comprises 7 subjects for a total of 61 single-source potentials.

### 2.2 Forward model

The forward model is a BEM model with realistic geometry. The model comprises three compartments and was set up using the function `make_bem_model` with *ico* set to 4, corresponding to a downsampling of the Freesurfer triangulations to 5,120 triangles; conductivities were automatically set to 0.3, 0.006 and 0.3 S/m, for the brain, skull and scalp compartments, respectively. The source space was built using 4,098 locations in each hemisphere, for a total of 8,196 available sources, with an average spacing of 4.9 mm.

### 2.3. Montages

We test inverse methods using four different sensor montages: the full montage contains 256 channels; then we repeatedly halve the number of channels going down to 128, then 64, and finally 32 channels. Note that in the case of 256 channels the effective number of channels is smaller than the nominal number, due to the removal of bad channels. On average the number of bad channels that were removed is 46 ± 23, mainly located over the neck and the chicks of the subjects, but, in some cases, also in the areas where the external part of intracranial electrodes were too dense to fit the hd-EEG net over the subject’s head.

### 2.4 Inverse methods

Source localization was carried out using ten different inverse methods. Nine of them are available as open source code within the MNE-Python package^2^ [42]: dipole fitting, dSPM [30], eLORETA [29], Gamma Map [32], Linearly Constrained beamformer [43], Mixed Norm Estimate [33], MNE [26], RAP-MUSIC [31], sLORETA[28]. In addition to these nine inverse algorithms, we also used SESAME [44, 34], a Bayesian multi-dipole modeling algorithm currently listed as a plug-in of MNE-Python. Our choice of working with MNE-Python was motivated by the following reasons: it contains the most used ESI methods; it contains the largest set of methods; last but not least, it is written in Python, a freely available programming language, while many alternative tools such as Fieldtrip [45] and Brainstorm [46] are available in Matlab. Along the paper, we will refer to each inverse method with its short name as listed in Table 1.

**Table 1:** Short name of inverse methods used in the study.

| Method | Short name |
| --- | --- |
| Dipole Fitting | DF |
| dSPM | DSPM |
| eLORETA | ELOR |
| Gamma Map | GM |
| Linearly Constrained beamformer | LCMV |
| Minimum Norm Estimate | MNE |
| Mixed Norm Estimate | MXNE |
| RAP-MUSIC | RAP |
| SESAME | SSM |
| sLORETA | SLOR |

In the analysis below we split the inverse methods in two classes according to a not completely standard classification: we call *distributed* methods those methods that are based on a distributed source model, and have no sparsity-encouraging penalty terms, i.e. MNE, dSPM, LCMV, SLOR, ELOR; we call *dipolar* methods both the methods based on strictly dipolar models such as DF, RAP and SSM, and also those methods based on a distributed source model but with a sparsity-encouraging penalty term, i.e MxNE and GM. This non-standard classification is motivated by the fact that MxNE and GM provide in output the estimated number of sources and the source locations, like purely dipolar methods do.

All methods, except DF and SSM, need a noise covariance matrix that was estimated from the pre-stimulus interval between −250 ms and −50 ms using the `compute_covariance` function in auto mode, in which cross-validation is used [47].

#### 2.4.1. Regularization parameters

All ESI methods under analysis, except DF and RAP, require the user to choose the value of one or more input parameters. In the following, we evaluate the performances of the methods when different values of parameters are used. For a fair comparison across methods, we let each parameter vary in the same interval and with the same values.

Seven ESI methods (all but DF, RAP and LCMV) require as input the SNR of the data, either directly or through the noise variance: for this parameter we test five different values ranging from 1, which has been recently shown to be a lower limit guaranteeing good accuracy of the reconstructions [15], to 5, that corresponds to extremely clean data. LCMV requires to set the regularization parameter λ for regularization of the covariance matrix: here we test five values, logarithmically spaced (see Table 2).

**Table 2:**
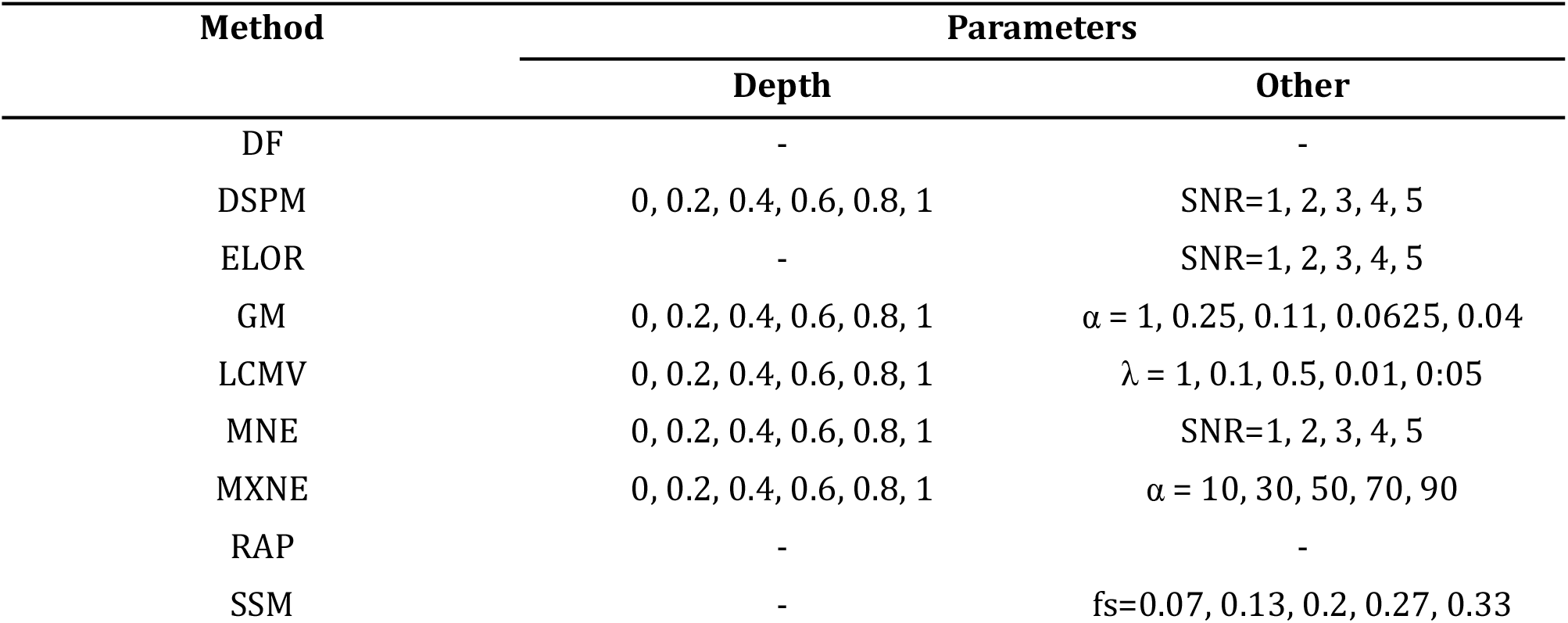
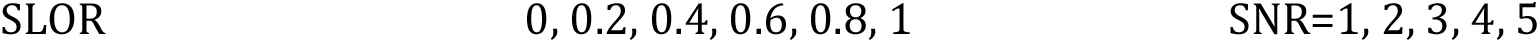
Parameters used for each inverse method.

| Method | Parameters |  |
| --- | --- | --- |
|  | Depth | Other |
| DF | - | - |
| DSPM | 0, 0.2, 0.4, 0.6, 0.8, 1 | SNR=1, 2, 3, 4, 5 |
| ELOR | - | SNR=1, 2, 3, 4, 5 |
| GM | 0, 0.2, 0.4, 0.6, 0.8, 1 | $\alpha = 1, 0.25, 0.11, 0.0625, 0.04$ |
| LCMV | 0, 0.2, 0.4, 0.6, 0.8, 1 | $\lambda = 1, 0.1, 0.5, 0.01, 0.05$ |
| MNE | 0, 0.2, 0.4, 0.6, 0.8, 1 | SNR=1, 2, 3, 4, 5 |
| MXNE | 0, 0.2, 0.4, 0.6, 0.8, 1 | $\alpha = 10, 30, 50, 70, 90$ |
| RAP | - | - |
| SSM | - | $f_s=0.07, 0.13, 0.2, 0.27, 0.33$ |

Finally, six methods also take in input a depth-weighting parameter, that aims to reduce the bias towards superficial sources: for this parameter we test five linearly spaced values between zero (no weight) and one. The different parameters we use and the corresponding values are reported in Table 2. We remark that both DF and RAP are parameter-free methods.

### 2.5. Performance evaluation

To quantify the source localization accuracy, we employ the Dipole Localization Error (DLE), which is defined as the distance between the estimated location and the putative dipole location, i.e. the medium point between the two electrodes in which current was fed. The estimated location is defined differently for distributed methods (DSPM, ELOR, LCMV, MNE, SLOR) and dipolar methods (DF, GM, MXNE, RAP, SSM). The distributed methods treat each time point independently; when applied to a time-series, they provide a (potentially) different intensity map/dipole location at each time step. For these methods we consider the solution at the peak latency and use the location corresponding to the peak intensity. For DF we consider the location of the equivalent dipole at the time point maximizing the goodness of fit in all analyzed time window. The remaining dipolar methods work natively with time-series, and provide one intensity map/dipole location(s) for the whole analysis window. For these methods we use the location of the estimated dipole applying the method to the window [−2; 2] ms; in case more than one dipole is estimated, we use the location of the estimated dipole with larger dipole moment. For all methods (except DF and RAP), we compute the DLE for the different parameters combination. In addition, for each method, we consider the mean solution defined as the average of the estimated dipole location over all parameters combination; for this mean dipole we compute the corresponding DLE with the putative dipole location.

Throughout the Results section, we will assess the differences between the performances of each and each other method for each montage with the use of the (non-parametric) Wilcoxon signed-rank test. We set the significance threshold at 0.05 and use Bonferroni correction for multiple comparisons.

## 3. Results

We present the results obtained by applying the ten inverse methods to all 61 sessions of the dataset. Figures 2 and 3 show an example of localization provided by the different inverse methods applied with five different values of a given input parameter, together with the exact location.

**Figure 2:**
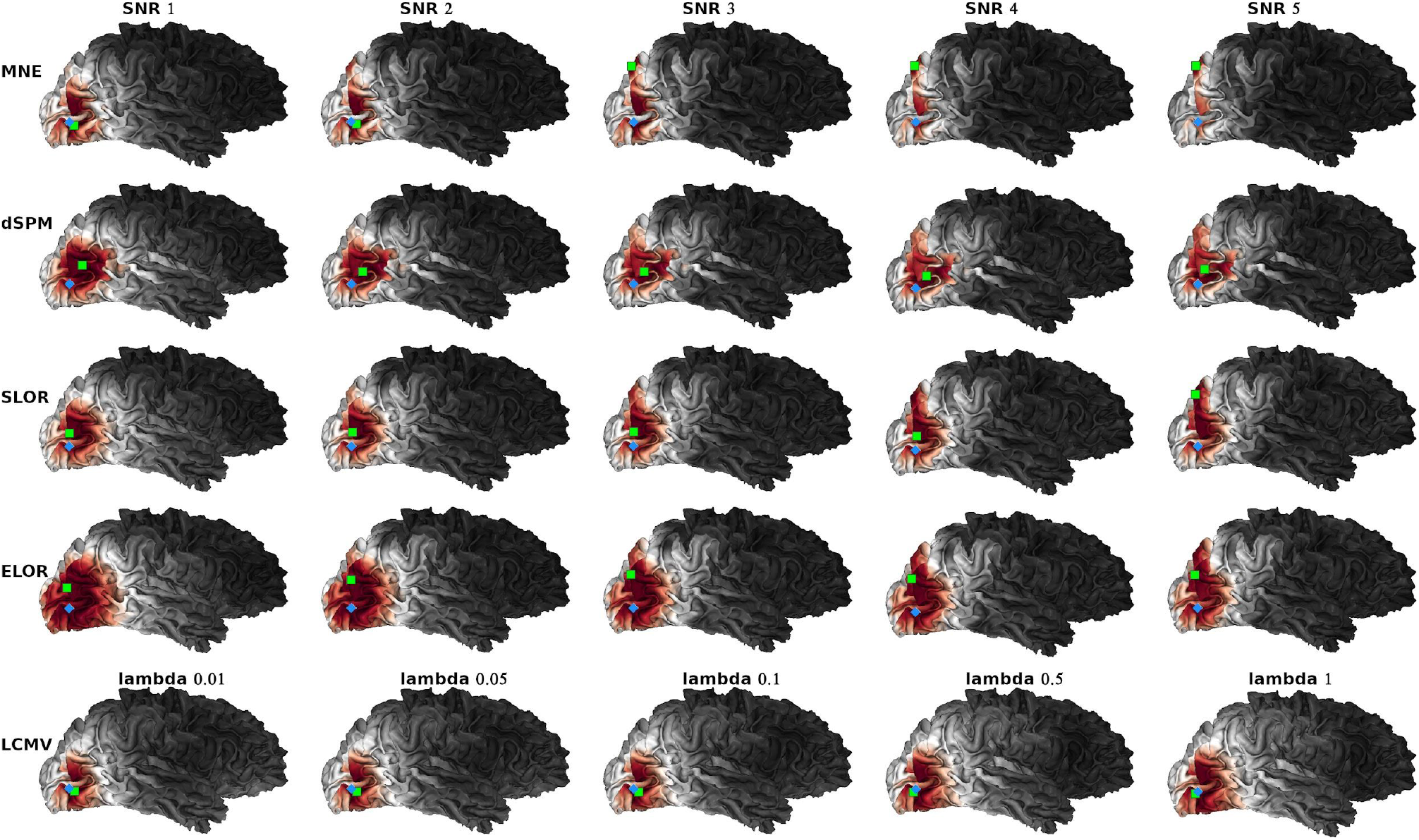
Example case: exact source location (light blue diamond), intensity map and estimated source location (green square) as obtained by the distributed methods (MNE. dSPM, SLOR, ELOR and LCMV) for five different values of the input parameter. The plot was done using the Visbrain ^3^suite [48].

**Figure 3:**
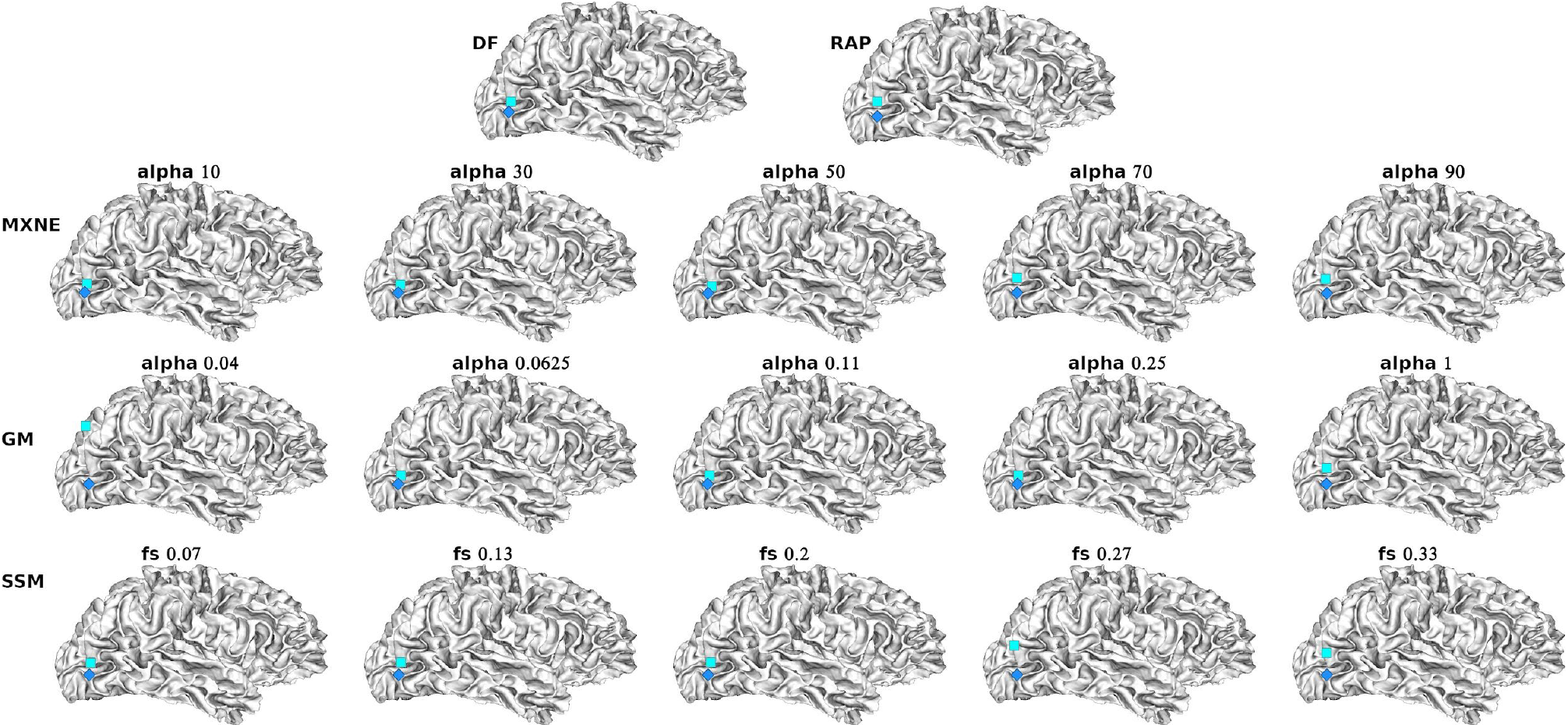
Example case: exact source location (light blue diamond) and estimated source location (cyan square) as obtained by the dipolar methods (DF, RAP, MXNE, GM and SSM.) For MXNE, GM and SSM the best solution is shown for five different values of the input parameter. The plot was done using the Visbrain suite [48].

### 3.1. Localization with the best combination of input parameters

Figure 4 contains the boxplots of minimum Dipole Localization Error (DLE) (in mm) computed for all methods and montages. For each method, for each session we consider the best solution across parameters (for more information on parameter values see Table 2), i.e. the one with the smallest DLE. Each method is coded by a specific color, and for each algorithm, from left to right, we show the boxplots obtained with 32, 64, 128, and 256 channels respectively. At the bottom of each boxplot we report its corresponding mean value (in mm). At a first glance, SSM and MXNE perform better than all other methods in all montages, while MNE seems the one with the worst performance.

**Figure 4:**
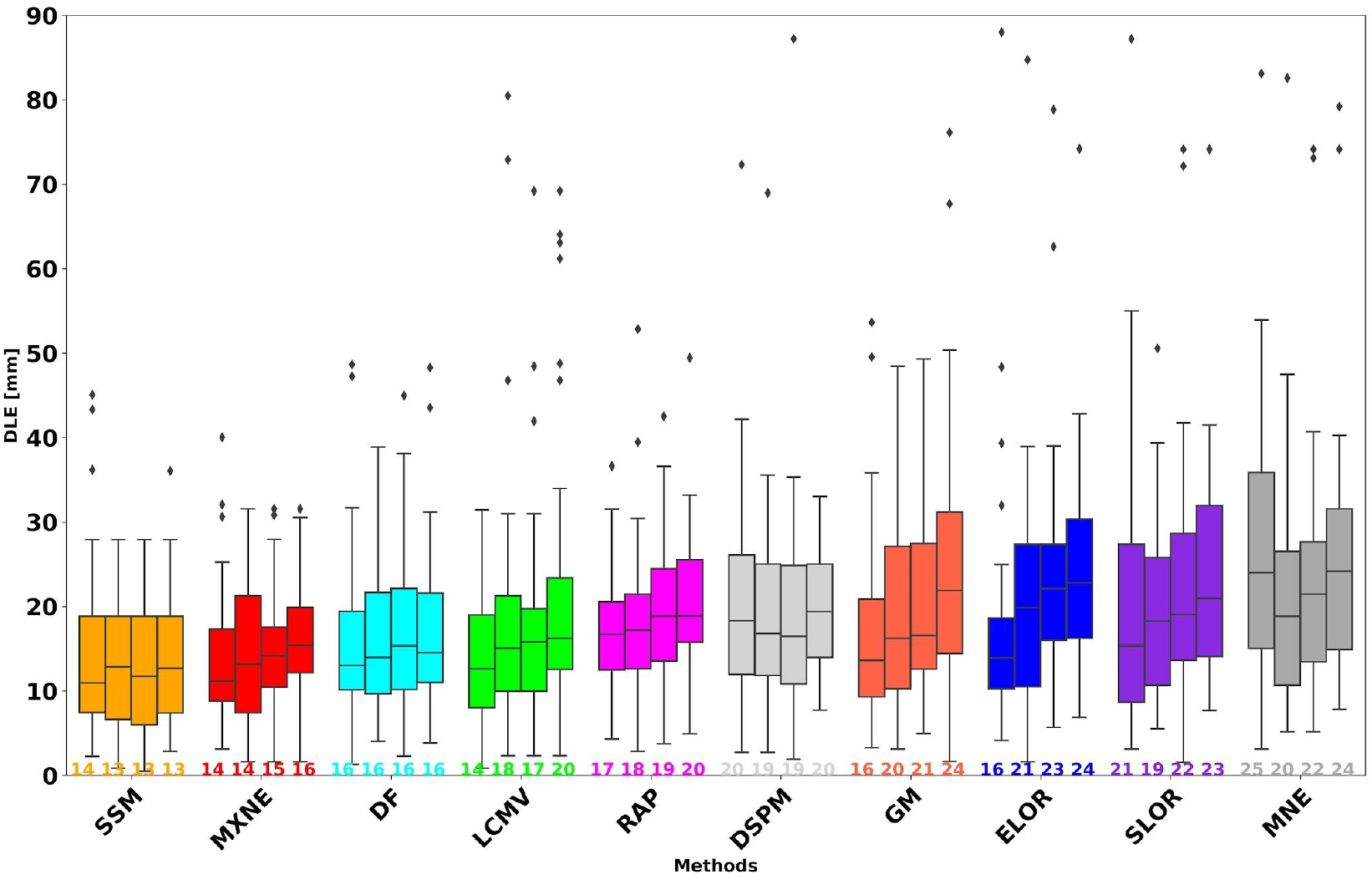
For each method, boxplot of minimum Dipole Localization Errors (DLE); from left to right, we show the boxplot obtained with 32 64, 128, and 256 channels, with (below) the mean value. ESI methods are ordered based on the overall behaviour.

In Figure 5, we report the results of the pairwise Wilcoxon signed-rank test assessing the difference between the performances of each and each other method when the best solution across parameters is considered. The results indicate that the best performances are obtained by: LCMV, MxNE and SSM with 32 channels; SSM and MXNE with 64 channels and DF, MxNE and SSM with 128 and 256 channels. Overall two dipolar methods, SSM and MxNE, substantially outperform the others.

**Figure 5:**
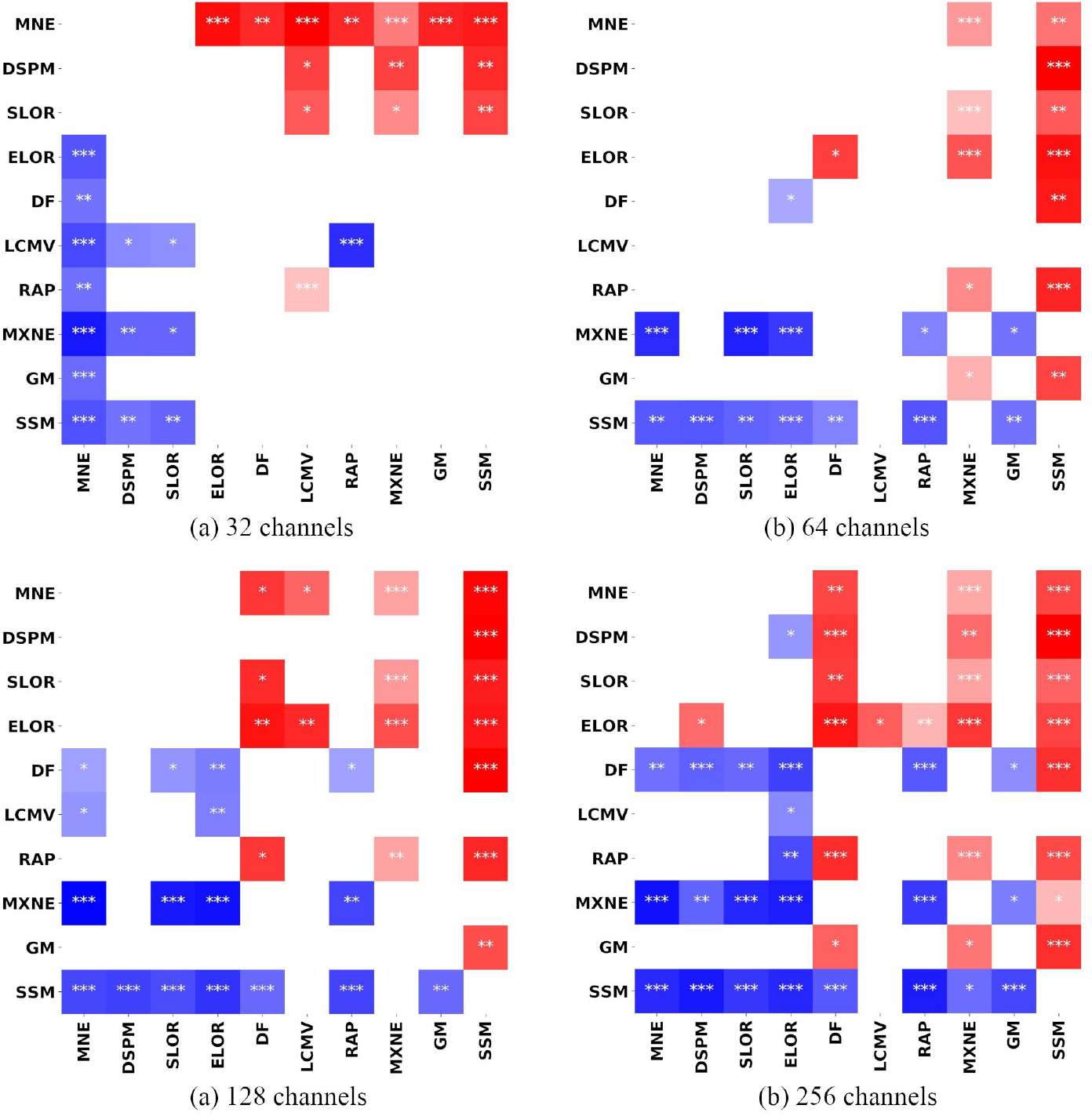
Significance of pairwise Wilcoxon tests between the DLE of each and each other ESI method, for the four different montages: 32 (first row left), 64 (first row right), 128 (second row left), and 256 channels (second row right). A red square indicates that the method listed in the corresponding row is significantly worse than the one listed in the corresponding column, while a blue square indicates that the method listed in the corresponding row is significantly better than the one listed in the corresponding column. The asterisks are related to the corrected p-value: * p < 0.05, ** p < 0.005, * * * p < 0.0005

Figure 6 shows, for each montage, the boxplot of the smallest DLE computed across all methods for each session; the blue boxplot represents the smallest DLE computed across all montages and methods. The median value for 32 channels is close to the one for 64, but the spread is smaller for 32 channels and the mean value is 8 mm with respect to the 9 mm of 64 sensors. In the case of 128 and 256 channels the global mean value is 8 and 9 mm respectively, while considering the smallest DLE across all montages and methods leads to a mean value of 6 mm and for more than half of the sessions we obtain a DLE < 5mm.

**Figure 6:**
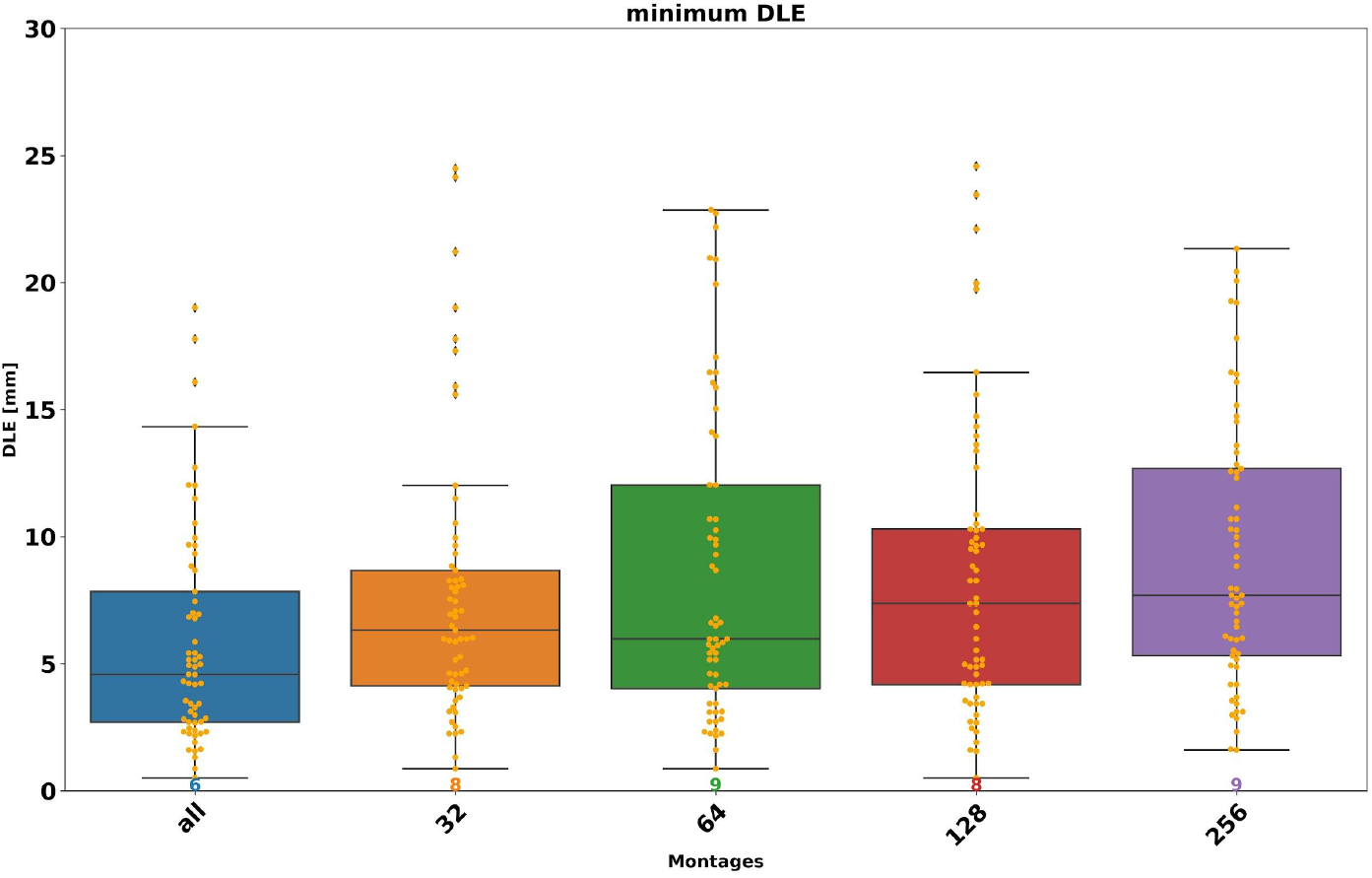
Minimum DLE over all methods for each montage. The blue boxplot represents the minimum of DLE across all montages.

In Figure 7 we report, for each method, the percentage of times the estimated location lies within 5 mm from the best solution obtained by the method. The method with the highest percentage is SSM, where for 256, 126 and 64 channels the smallest DLE is obtained more than 60% of times, while by decreasing the number of channels the minimum is reached less times (51%). The smallest DLE is reached for almost half of the sessions by LCMV (256, 128 and 32 channels), MXNE (64 and 32 channels) and DF (256 channels). ELOR and SLOR reach the minimum more times when fewer channels are considered, MNE and DSPM with 64 channels and RAP and GM with 128 channels.

**Figure 7:**
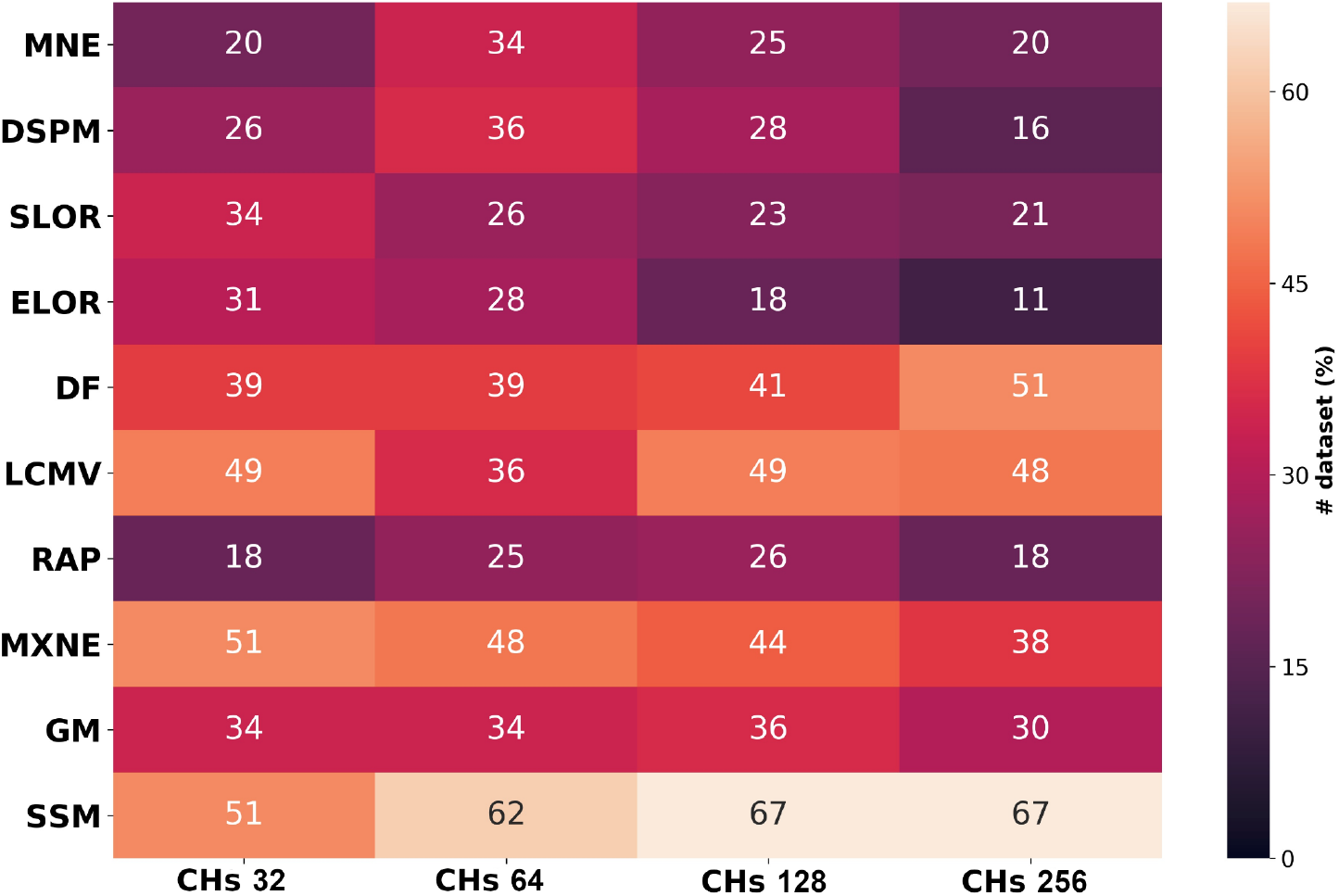
Number of times (%) in which the minimum DLE for each montage is reached by each method by using a tolerance δ = 5 mm.

Finally, we tested pairwise each method and each montage. We obtained significantly different values for many of the distributed methods (dSPM, ELOR, MNE, SLOR) and GM and RAP among the dipolar methods. The significant p-values of these statistical tests are reported in Table 3.

**Table 3:** For each method, significance of pairwise Wilcoxon signed-rank tests between the DLE of each and each other montage. We report the corrected p-values.

| Method | Channels | p-value |
| --- | --- | --- |
| MNE | 256 > 128 | $3.2 \times 10^{-2}$ |
| | 256 > 64 | $2.8 \times 10^{-4}$ |
| | 128 > 64 | $1.9 \times 10^{-2}$ |
| | 64 < 32 | $1.1 \times 10^{-3}$ |
| DSPM | 256 > 128 | $1.0 \times 10^{-2}$ |
| SLOR | 256 > 64 | $6.2 \times 10^{-4}$ |
| | 128 > 64 | $1.3 \times 10^{-3}$ |
| ELOR | 256 > 128 | $1.6 \times 10^{-2}$ |
| | 256 > 64 | $7.6 \times 10^{-3}$ |
| | 256 > 32 | $8.9 \times 10^{-7}$ |
| | 128 > 64 | $1.5 \times 10^{-2}$ |
| | 128 > 32 | $1.1 \times 10^{-5}$ |
| | 64 > 32 | $2.8 \times 10^{-3}$ |
| RAP | 256 > 64 | $6.4 \times 10^{-3}$ |
| | 256 > 32 | $4.7 \times 10^{-3}$ |
| GM | 256 > 128 | $2.4 \times 10^{-2}$ |
| | 256 > 64 | $7.8 \times 10^{-4}$ |
| | 256 > 32 | $4.1 \times 10^{-5}$ |
| | 128 > 32 | $6.7 \times 10^{-3}$ |
| | 64 > 32 | $3.7 \times 10^{-2}$ |

### 3.2. Localization accuracy and input parameters

To study the influence of input parameters on localization accuracy, for each montage and method, given a session, we compute the solutions corresponding to different combinations of parameter(s) values; we then consider the mean dipole and its DLE, as well as the standard deviation with respect to the mean dipole as∷

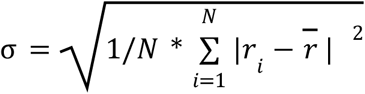

where *N* is the number of parameters combination (see Table 2), *r_i_* is the location of the dipole estimated by the ith set of parameters and 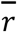 is the location of the mean dipole computed across all N parameters combinations.

In Figure 8 we report the DLE (top) and the standard deviation (bottom) computed with respect to the mean dipole. By visually comparing the results in Figure 4 and 8(a), we observe that most of the methods behave in a similar way, and the difference between the DLE of the global mean and that of the best solution ranges between 2 and 7 mm. We remark that DF and RAP do not depend on parameters, thus the corresponding boxplots are the same as in Fig. 4. The only notable difference is for LCMV that performs worse, probably due to the great variability of the estimated dipoles for the different combination of the parameters. This behaviour emerges from Figure 8(b) that gives us information on the variability of the location of the estimated dipole across all parameters combinations. Among distributed methods, DSPM has a very low standard deviation, while among the dipolar methods, SSM is the one with less spread around the mean dipole. Conversely, LCMV for all the montages except 32 channels presents a very high variability of the solutions for the different input parameters.

**Figure 8:**
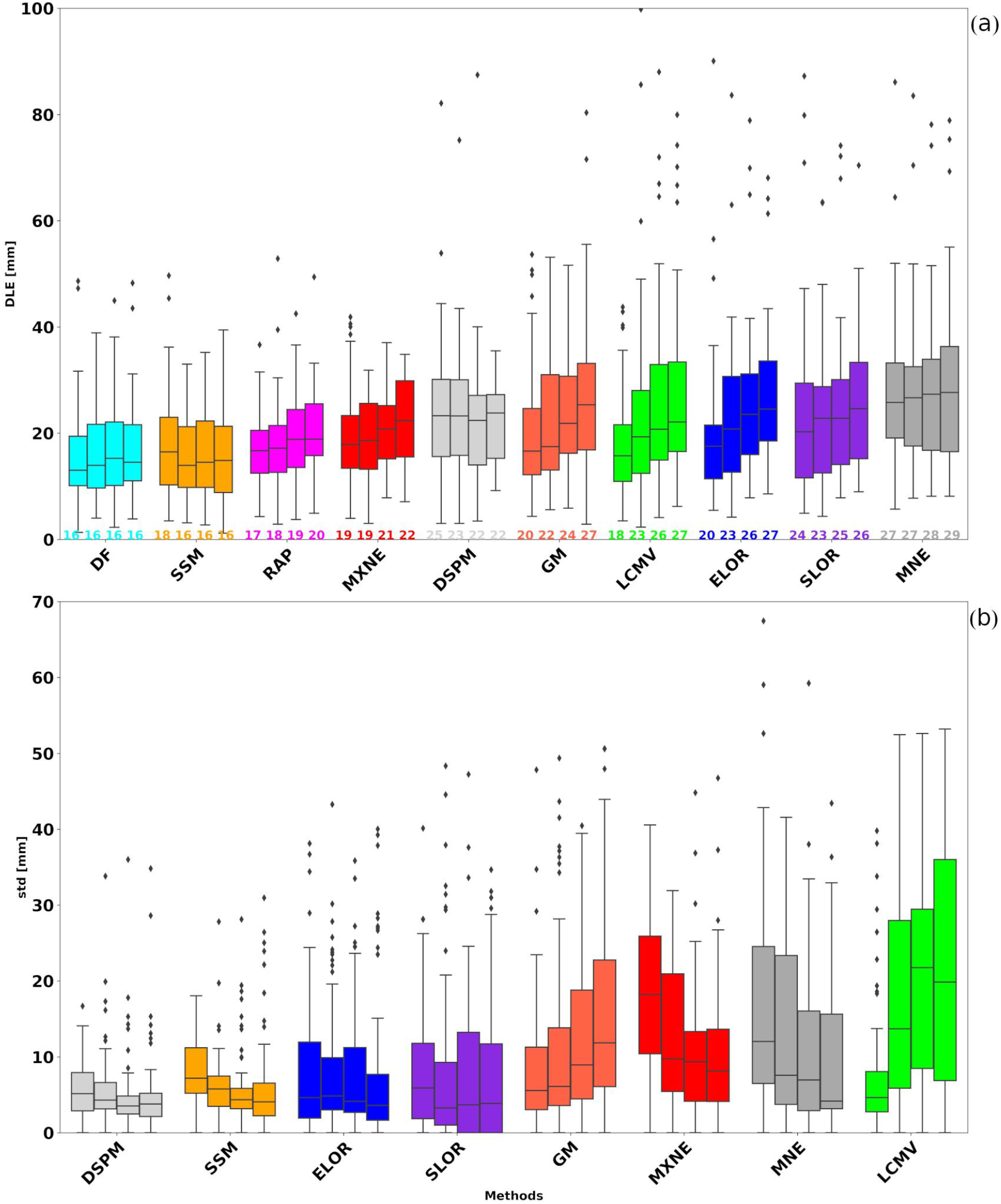
From left to right for each method we show results with 32, 64, 128, and 256 channels. (a) Dipole Localization Error (DLE) computed by using the mean dipole over all parameter combinations; below, the mean value. (b) Standard deviation of the DLE across all parameter combinations.

Figure 9 reports the results of the statistical tests of each and each other method for each montage with the use of the (nonparametric) Wilcoxon signed-rank test. By comparing the results obtained with the mean dipole and the ones of Fig. 5, we can see that the statistical difference between different inverse methods is more significant when the mean dipole is used, mainly for a high number of channels: with 256, 128 and 64 channels we have similar patterns with the best performance obtained by DF, MxNE, RAP and SSM; with 32 channels MNE performs worse than all other methods.

**Figure 9:**
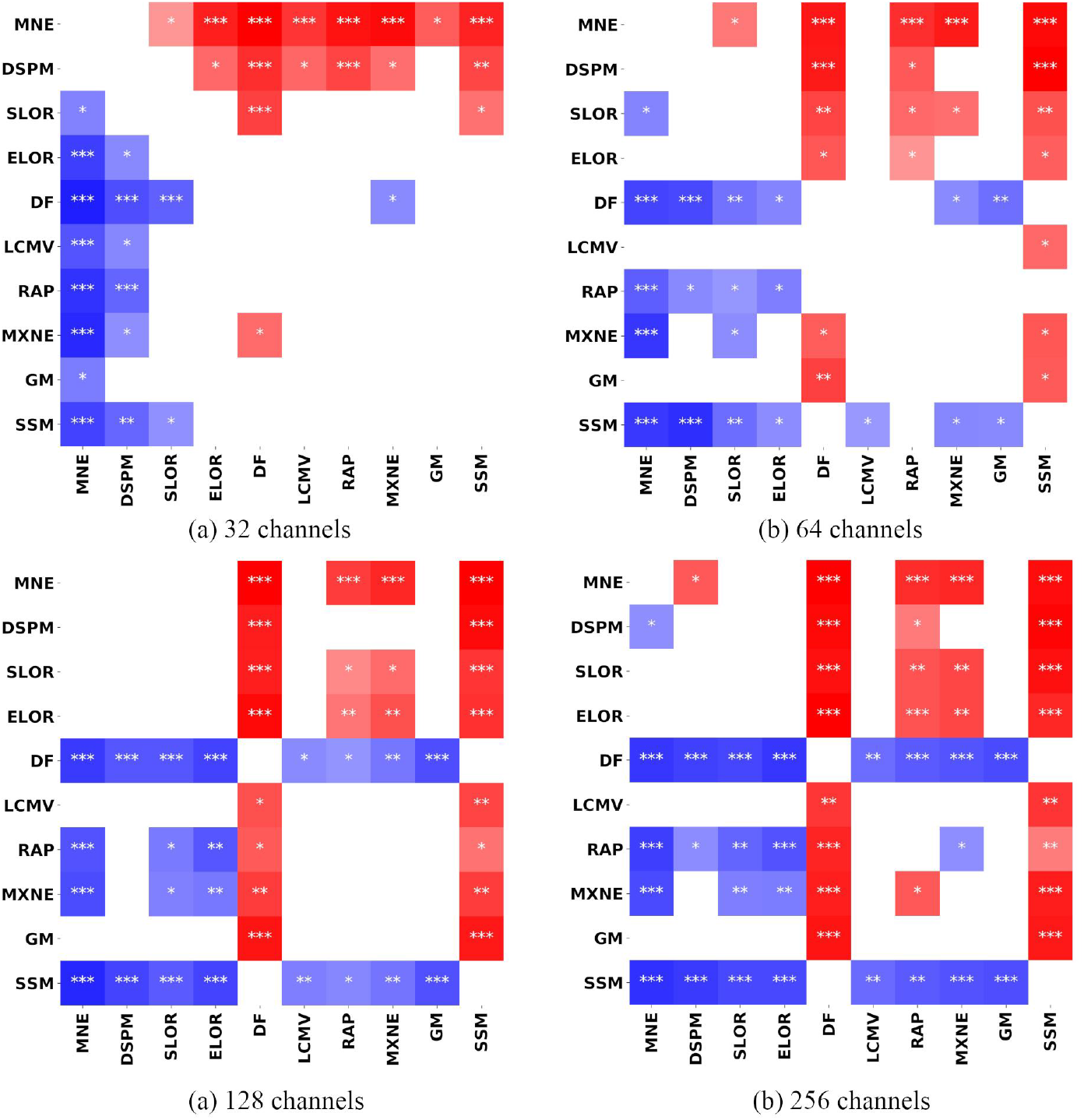
Significance of pairwise Wilcoxon tests between the DLE of the mean dipole of each and each other inverse method, for the four different montages: 32 (first row left), 64 (first row right), 128 (second row left), and 256 channels (second row right). A red square indicates that the method on row is significantly worse than the corresponding method on column, while a blue square indicates that the method on row is significantly better than the corresponding one on column. The asterisks are related to the p-value: * p < 0.05, ** p < 0.005, * * * p < 0.0005

Figure 10 gives an insight into the influence of parameters on the solution of the different inverse methods. For each method we report, for each combination of input parameter values, the percentage of times the estimated location lies within 5 mm from the best solution obtained by the method across all parameter combinations. Among distributed methods, MNE and MxNE appear to be the most sensitive to the depth parameter, while other distributed methods seem to be only influenced by the SNR parameter; for LCMV it seems there is only a negative trend when no depth weighting is performed (i.e. depth=0). For MXNE a value of alpha between 30 and 70 together with a value for depth equal to 1 gives the best result. For GM as for the distributed methods the most important parameter is the noise variance while SSM shows better performances for smaller values of the parameter, corresponding to higher SNR.

**Figure 10:**
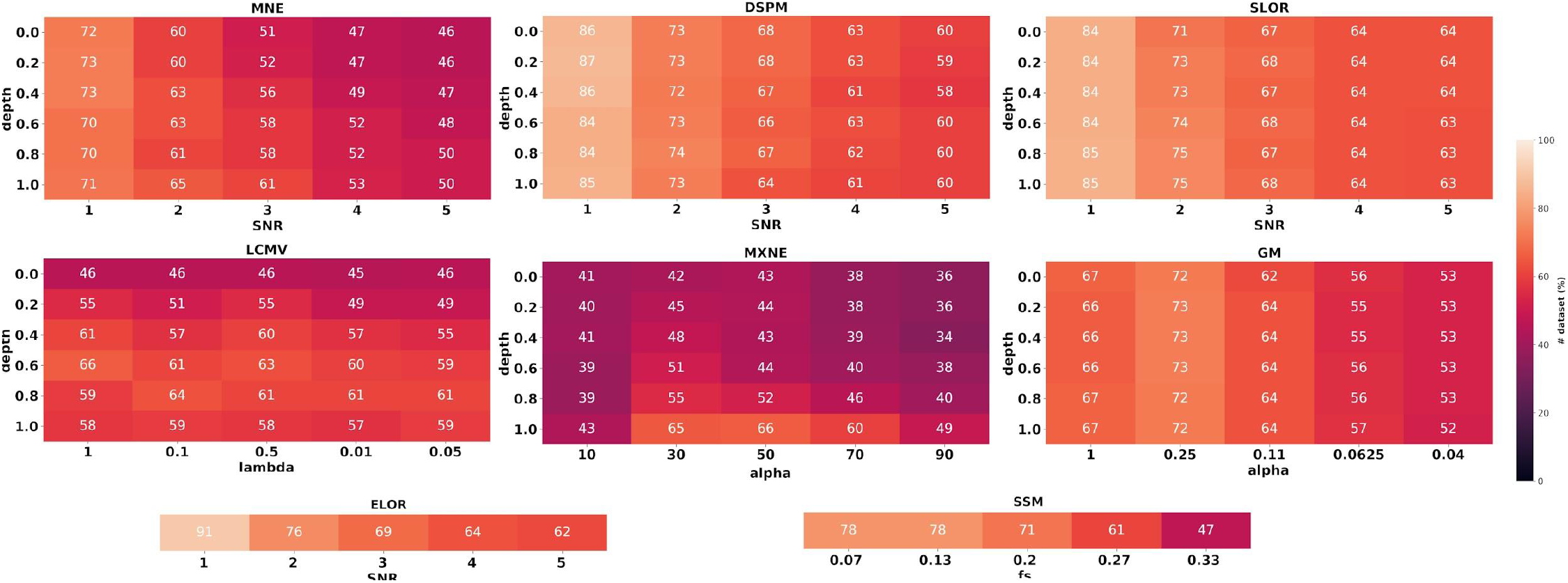
Percentage of times (%) a specific combination of parameter values reaches within 5 mm from the best solution obtained by the method across all parameter combinations.

## 4. Discussion

The aim of this work was to evaluate and compare *in vivo* the localization accuracy of a relatively large set of ESI methods, under the hypothesis that the neural generators are focal sources and substantially dipolar.

### 4.1. Localization with the best combination of input parameters

When using the best combination of input parameters the results are encouraging: the best solution across methods is within 1 cm from the true source with very high probability, and several methods provide average reconstruction errors around 1 cm, with about 75% of cases falling within 2 cm. It is important to remark that in this study the source reconstruction procedure was completely automated: after application of the ESI method, only the stronger source was retained, leading sometimes to large errors. While in routine analysis it may be difficult to select the optimal combination of parameters, the user may leverage on prior knowledge and sometimes exclude some of the reconstructed sources, thus effectively reducing the localization error estimated in our study.

Interestingly, we observed substantial and significant variability between methods, with MNE being the least accurate and SSM the most accurate. As expected, dipolar methods provided better results than distributed methods. In this respect, it may be worth to observe that there is an important difference between the distributed and the dipolar methods considered here: indeed SSM, RAP, GM and MxNE make use of the whole time series, while MNE, SLOR, ELOR, dSPM and LCMV work on a single time point and might be more affected by noise; on the other hand, the SNR of the data is quite high. We also notice that, pleasantly, newer methods tend to outperform older ones, confirming that there is progress in the field.

We reckon the residual error we observe is most likely due to the combined effect of bias introduced by the ESI method, and forward modeling error. In this respect, we recall that the single pulse stimulation is a squared wave lasting 1 millisecond, and therefore contains very high frequency components: future studies might be devoted to investigating whether the quasi-static approximation is still valid under these circumstances.

### 4.2. Sensitivity to input parameters

In general, input parameters do impact localization accuracy of ESI methods quite substantially. However the actual impact of the input parameter is larger for some methods and smaller for others. We quantified this variability by computing the standard deviation of the DLE across parameter combinations. Notice that, a priori, one would expect that methods with high variability provide a better best result, because more variability implies more chances of getting closer to the true solution at least once. Indeed, by comparing Figure 8 and 4 we see that this holds true for MxNE and LCMV, that score quite well in terms of best solution and relatively high in terms of variability. There are also exceptions to this rule: SSM has one of the lower standard deviations while being the most accurate with the best solution; MNE, on the other hand, has high variability but also high localization error. Hence, some methods are more robust against a mis-specification of the input parameters, and some are less.

We also observed two unexpected results. First, for distributed methods we observed the best performances in correspondence of SNR=1, i.e. the smallest value of the SNR parameter: even though the reconstructions become more widespread, the peak gets closer to the true source. This result is puzzling because SNR=1 corresponds in principle to very noisy signals, while the input data are quite clean; on the other hand, it finds partial confirmation in [49] where authors use SNR=3 for MEG data and SNR=1 for EEG data. We speculate this fact might be due to forward modeling errors, related to the volume conduction problem, that reduce the effective SNR of otherwise clean EEG data; in any case, more investigations are needed to clarify this point. Second, despite the presence of both deep and superficial sources in the dataset, the depth weighting parameter appears to have little or no impact for half of the tested methods, namely dSPM, SLOR and GM; in MNE and MxNE, on the other hand, the largest tested value provided the best results, while LCMV seems to prefer intermediate values.

As a side note, it is interesting to compare the low standard deviation of dSPM with the high standard deviation of MNE. Indeed, we recall that the dSPM solution is obtained from the MNE solution through a “noise normalization” procedure. Apparently, such noise normalization contributes a little to reducing localization error but mostly to reduce the dependence on the input parameters.

Overall, our results clearly point out that objective and reliable criteria for choosing the parameters are needed.

### 4.3. Impact of the montage

In our study, the best localization accuracy was often obtained with 32 channels, and we observed no major differences when using denser montages; in fact, for some methods we found significant differences in the DLE obtained with different montages, and the lower DLE was almost systematically associated to the lower density montage. While this result is certainly unexpected, there are few considerations that can help to make it less counter intuitive than it appears at first.

Indeed, there are two specific features of the analyzed data set that make 32 channels an enough number of channels: (i) the dipolar nature of the source we are looking for and (ii) the high SNR of the data. Indeed, localizing a single current dipole amounts to estimating 6 parameters: 32 channels, corresponding to 32 equations or constraints, are more than enough provided that their data are good enough [50]. And the data are good enough, because the spatial distribution of the 32 channels covers the whole head, and because the SNR of the data is very high. Therefore, under our experimental conditions, we do not expect a substantial gain in localization accuracy when adding more channels. This result is in agreement with literature on the topic [51, 52].

From a mathematical perspective, denser montages correspond to taller leadfield matrices featuring larger condition numbers, i.e. the problem becomes more ill-conditioned and more regularization might be required. As a partial confirmation, significant differences between montages were observed almost exclusively for distributed methods, that are expected to suffer more from an ill-conditioned leadfield; the best DLE of SSM, DF, MxNE and LCMV showed no significant dependence on the montage.

Finally, we notice that several methods show reduced variability with respect to the input parameter when denser montages are used: in this respect, denser montages do provide better results in terms of increased stability.

In conclusion, our results suggest that 32 channels are enough to reconstruct a focal source from high-SNR data; this might include single time points with one strong source, but also single topographies obtained by ICA, or specific frequencies. More complex configurations (or more noisy data sets), on the other hand, are likely to benefit from additional sensors.

### 4.4. Comparison with previous works

Several comparisons between ESI methods have been performed in recent years.

First of all, the same dataset used here was used in [25], where an exemplar analysis with three ESI methods (MNE, ELOR and DSPM) was performed. Here, we considerably extended the comparison to include also more recent methods: noteworthy, we observed that recent methods such as SSM and MxNE do outperform older ones, hereby confirming recent results [19, 15]. In addition, we studied the impact of regularization and depth weighting parameters more in detail, highlighting similarities and differences between different methods.

Another study that relates quite directly to our current study is that in [19], where the authors compared retrospectively SSM, RAP and wMNE with the results of an ECD analysis on epileptic subjects; indeed, also in this case the reference source is a point source, even though in this case it is the product of a former analysis and not a true source; anyway, the authors find MNE to be the least accurate and SSM the most accurate.

There is also an increasing number of studies that find different methods have substantially similar, good performances. In [53] the authors compare 5 ESI methods on ictal EEG data, and find a general agreement between methods, with MNE being the least accurate; although this result is not statistically significant, it is a partial confirmation of our result. In [20] the authors compared DSPM, MNE, SLOR and cMEM [54] (not tested here) in a clinical scenario, and found excellent performances for all of them. Their results show sLOR was slightly but significantly better than dSPM, while in our findings SLOR was at times better than MNE but never better than dSPM. In [55] the authors study the accuracy of dipolar methods (ECD, MUSIC), imaging methods (MNE, sLORETA, SWARM) and different implementation of SAM beamformer as compared to intracranial EEG (iEEG) and the resection areas in a large cohort of pediatric patients with intractable epilepsy. The accuracy of all these methods is relatively similar when compared to the ground truth. The concordance or discordance of MUSIC with iEEG was the best predictor of long-term seizure outcome. In [56] the authors recently compared interictal MEG spikes using MUSIC, SAM(g2), and sLORETA to interictal discharges recorded with iEEG. It was reported that these three MEG methods showed similar concordance with iEEG but differed depending on the brain region in which the spike was located.

In other comparisons, the authors use dataset where more widespread activations are present. In [17] the authors study what are the best conditions for locating the epileptogenic zone with High Resolution EEG and compare five different methods; their results highlight that distributed methods are more appropriate to localize a widespread epileptogenic zone than a focal one, and that ictal spikes with focal scalp electric field are better localized by dipolar methods (ECD and MUSIC). This last finding is in line with our results. In [57] the authors used MNE, eLOR and LCMV for source reconstruction and connectivity estimation from resting state data, and found relative agreement in source localizations, more than in connectivity estimation.

Most of these studies feature three important differences with respect to the one reported here. First of all, the definition of true source was necessarily more vague and less accurate than here. Second, the data were analyzed by expert users who almost certainly had expectations, and could tune each method to provide a coherent picture; here, instead, each method was applied independently, and in an automated fashion. Third, none of these studies makes explicit reference to the setting of the input parameters.

## 5. Conclusions

In this study we investigated in vivo the spatial accuracy of ESI methods, and its dependence on the input parameter(s), when the generators are focal. Our data show good levels of accuracy of ESI techniques, with the best solution across parameters and across methods within 1 cm from the true source. This is true also when ESI is applied to “conventional” (32 channels) rather than dense (64, 128, 256 channels) EEG recordings. While all tested methods provided reasonable performances with the optimal parameter value(s), we did observe substantial differences between methods: recent dipolar methods, particularly SSM, provide significantly better results than older distributed methods such as MNE, both in terms of a higher accuracy with the optimal parameter choice, and a lower sensitivity to the value of the input parameter. We also observed negligible impact of depth weighting for SLOR, dSPM and GM, and a general preference for lower values of the SNR in all distributed methods, a result that finds partial support in the literature.

## Acknowledgments

This research has received funding from the European Union’s Horizon 2020 Framework Programme for Research and Innovation under the Specific Grant Agreements No. 785907 and No. 945539 (Human Brain Project SGA2 and SGA3). Annalisa Rubino and Lino Nobili received funding from the Italian Ministry of Health, Targeted ResearchGrant No. RF-2010-2319316. This work was developed within the framework of the DINOGMI Department of Excellence of MIUR 2018-2022 (law 232/2016).

## Footnotes

2 https://mne.tools

3 http://visbrain.org/

